# SciPipe - A workflow library for agile development of complex and dynamic bioinformatics pipelines

**DOI:** 10.1101/380808

**Authors:** Samuel Lampa, Martin Dahlö, Jonathan Alvarsson, Ola Spjuth

## Abstract

**Background:** The complex nature of biological data has driven the development of specialized software tools. Scientific workflow management systems simplify the assembly of such tools into pipelines, assist with job automation and aid reproducibility of analyses. Many contemporary workflow tools are specialized and not designed for highly complex workflows, such as with nested loops, dynamic scheduling and parametrization, which is common in e.g. machine learning.

**Findings:** SciPipe is a workflow programming library implemented in the programming language Go, for managing complex and dynamic pipelines in bioinformatics, cheminformatics and other fields. SciPipe helps in particular with workflow constructs common in machine learning, such as extensive branching, parameter sweeps and dynamic scheduling and parametrization of downstream tasks. SciPipe builds on Flow-based programming principles to support agile development of workflows based on a library of self-contained, reusable components. It supports running subsets of workflows for improved iterative development, and provides a data-centric audit logging feature that saves a full audit trace for every output file of a workflow, which can be converted to other formats such as HTML, TeX and PDF on-demand. The utility of SciPipe is demonstrated with a machine learning pipeline, a genomics, and a transcriptomics pipeline.

**Conclusions:** SciPipe provides a solution for agile development of complex and dynamic pipelines, espe-cially in machine leaning, through a flexible programming API suitable for scientists used to programming or scripting.

## Findings

Driven by the highly complex and heterogeneous nature of biological data [1, 2], computational biology is characterized by an extensive ecosystem of command-line tools, each specialized on one or a few of the many aspects of biological data. Because of their specialized nature these tools generally need to be assembled into sequences of processing steps, often called “pipelines”, to produce meaningful results from raw data. Due to the increasingly large sizes of biological data sets [3, 4], such pipelines often require integration with High-Performance Computing (HPC) infrastructures or cloud computing resources to complete in an acceptable time. This has created a need for tools to coordinate the execution of such pipelines in an efficient, robust and reproducible manner. This coordination can in principle be done with simple scripts in languages like Bash, Python or Perl, but such scripts can quickly become fragile.

When the number of tasks becomes sufficiently large, and the execution time sufficiently long, the risk for failures during the execution of such scripts increases almost linearly with time, and simple scripts are not a good strategy for when large jobs need to be restarted from a failure. They lack the ability to distinguish between finished and half-finished files, and can not by default detect if intermediate output files are already created and can be reused to save computing time and resources. These limits with simple scripts calls for a strategy with a higher level of automation. This need is addressed by a class of software commonly referred to as “scientific workflow management systems” or simply “workflow tools”. Through a more automated way of handling the execution, workflow tools can improve the robustness, reproducibility and understandability of computational analyses. In concrete terms, workflow tools provide means for handling atomic writes (making sure finished and half-finished files can be separated after a crashed or stopped workflow), caching of intermediate results, distribution of tasks to the available computing resources and automatically keeping or generating records of exactly what was run, to make analyses reproducible.

It is widely agreed upon that workflow tools generally make it easier to develop automated, reproducible and fault-tolerant pipelines, although many challenges and potential areas for improvement still do exist with existing tools [5]. This has made scientific workflow systems a highly active area of research. Numerous workflow tools have been developed and many new ones are continuously being developed.

The workflow tools developed differ quite widely in terms of how workflows are being defined and what features are included out-of-the box. This probably reflects the fact that different types of workflow tools can be suited for different categories of users and use cases. Graphical tools like Galaxy [6, 7, 8] and Yabi [9] provide easy to use environments especially well-suited for scientists without scripting experience. Text-based tools like Snakemake [10], Nextflow [11], BPipe [12], Cuneiform [13] and Pachyderm [14] on the other hand, are implemented as Domain Specific Languages (DSLs), that can often provide a higher level of flexibility, at the expense of the ease of use of a graphical user interface. They can thus be well suited for “power users” with experience in scripting or programming.

Even more power and flexibility can be gained from workflow tools implemented as programming libraries, which provide their functionality through a programming API accessed from an existing programming language such as Python, Perl or Bash. By implementing the API in an existing language, users get access to the full power of the implementation language when writing workflows, as well as the existing tooling around the language. One example of a workflow system implemented in this way is Luigi [15].

As reported in [5], although many users find important benefits in using workflow tools, many also experience limitations and challenges with existing workflow tools, especially regarding the ability to express complex workflow constructs such as branching and iteration, as well as limitations in terms of audit logging and reproducibility. Below we will briefly review a few of existing, popular systems, and highlight areas where we found that the development of a new approach and tool was desirable for use cases that includes very complex workflow constructs.

Firstly, graphical tools like Galaxy and Yabi, although being easy-to-use even for users without programming experience, is often perceived to be limited in their flexibility due to the need to install and run a web server in order to use them, which is not always permitted, or practical, on HPC systems.

Text-based tools implemented as DSLs, such as Snakemake, Nextflow, BPipe, Pachyderm and Cuneiform do not have this limitation, but have other characteristics which might be problematic for for complex workflows in some cases.

For example, Snakemake is *dependent* on file naming strategies for defining dependencies, which can in some situations be limiting, and also uses a “pull-based” scheduling strategy (the workflow is invoked by asking for a specific output file, where after all tasks required for reproducing the file will be executed). While this makes it very easy to reproduce specific files, it can make the system hard to use for workflows involving complex constructs such as nested parameter sweeps and cross-validation fold generation, where the final file names might be hard if at all possible to foresee. Snakemake also performs scheduling and execution of the workflow graph in separate stages, which means that it does not support dynamic scheduling.

Dynamic scheduling, which basically means on-line scheduling during the workflow execution [16], is useful both where the number of tasks is unknown before the workflow is executed, and where a task needs to be scheduled with a parameter value obtained during the workflow execution. An example of the former is reading row by row from a database, splitting a file of unknown size into chunks, or processing a continuous stream of data from an external process such as an automated laboratory instrument. An example of the latter is training a machine learning model with hyperparameters obtained from a parameter optimization step prior to the final training step.

BPipe constitutes a sort of middle-ground in terms of dynamic scheduling. It supports dynamic decisions of what to run by allowing execution-time logic inside pipeline stages, as long as the structure of the workflow does not need to change. Dynamic change of the workflow structure can be important in workflows for machine learning though, e.g. if parametrizing the number of folds in a cross-validation based on a value calculated during the workflow run, such as dataset size.

Nextflow, has push-based scheduling and supports full dynamic scheduling via the dataflow paradigm, does not have this limitation. It does not, however, support creating a library of reusable workflow components, because of its use of dataflow variables shared across component definitions, as this requires processes and the workflow dependency graph to be defined together.

Pachyderm is a container-based workflow system which uses a JSON and YAML-based DSL to define pipelines. It has a set of innovative features including a version-controlled data management component with Git-like semantics and support for triggering of pipelines based on data updates, among others. These in combination can provide some aspects of dynamic scheduling. On the other hand, the more static nature of the JSON/YAML-based DSL might not be optimal for really complex setups such as creating loops or branches based on parameter values obtained during the execution of the workflow. The requirement of Pachyderm to be run on a Kubernetes [17] cluster can also make it less suitable for some academic environments where ability to run pipelines also on traditional HPC clusters is required. On the other hand, because of the easy incorporation of existing tools, it is possible to provide such more complex behavior by including a more dynamic workflow tool as a workflow step in Pachyderm instead. We thus primarily see Pachyderm as a complement to other light-weight workflow systems, rather necessarily than an alternative.

The usefulness of such an approach where an over-arching frameworks provide primarily an orchestration role while calling out to other systems for the actual workflows, is demonstrated by the Arteria project [18]. Arteria builds on the event-based StackStorm framework to allow triggering of external workflows based on any type of event, providing a flexible automation framework for sequencing core facilities.

Going back to traditional workflow systems, Cuneiform takes a different approach compared to most workflow tools by wrapping shell commands in functions in a flexible functional language (described in [19]), which allows leveraging common benefits in functional programming languages such as side-effect free functions, to define workflows. It also leverages the distributed programming capabilities of the Erlang Virtual Machine (EVM), to provide automatic distribution of workloads. It is still a new, domain specific language though, which means that tooling and editor support might not be as extensive as for an established programming language.

Luigi is a workflow library developed by Spotify, which provides a high degree of flexibility due to its implementation as a programming library, Python. For example, the programming API exposes full control over file name generation. Luigi also provides integration with many Big Data systems such as Hadoop and Spark, and cloud-centric storage systems like HDFS and S3.

SciLuigi [20] is a wrapper library for Luigi, previously developed by the authors, which introduces a number of benefits for scientific workflows by leveraging selected principles from Flow-based programming (FBP) (named ports and separate network definition) to achieve an API that makes iteratively changing the workflow connectivity easier than in vanilla Luigi.

While Luigi and SciLuigi were shown to be a helpful solution for complex workflows in drug discovery, they also have a number of limitations for highly complex and dynamic workflows. Firstly, since Python is an untyped, interpreted language, certain software bugs are discovered only far into a workflow run, rather than while compiling the program. Secondly, the fact that Luigi creates separate processes for each worker, which communicate with the central Luigi scheduler via HTTP requests over the network, can lead to robustness problems when going over a certain number of workers (around 64 in the authors’ experience) leading to HTTP connection time-outs.

The mentioned limitations for complex workflows in existing tools is the background and motivation for developing the SciPipe library.

### The SciPipe workflow library

SciPipe is a workflow library based on Flow-Based Programming principles, implemented as a library in the Go programming language. The library is freely available as open source on GitHub [21]. All releases of GitHub are also archived on Zenodo [22]. Similarly to Nextflow, SciPipe leverages the dataflow paradigm to achieve dynamic scheduling of tasks based on input data, allowing many workflow constructs not easily coded in many other tools.

Combined with design principles from Flow-based programming such as separate network definition and named ports bound to processes, this has resulted in a productive and discoverable API that enables agile authoring of complex and dynamic workflows. The fact that the workflow network is defined separately from processes, enables building workflows based on a library of reusable components, although the creation of ad-hoc shell-command based components is also supported.

SciPipe provides a provenance tracking feature that creates one audit log per output file, rather than only one for the whole workflow run. This means that it is always easy to verify exactly how each output of a workflow was created.

SciPipe also provides a few features which are not very common among existing tools, or which are not commonly occurring together in one system. These include support for streaming via Unix named pipes, ability to run push-based workflows up to a specific stage of the workflow, and flexible support for file naming of intermediate data files generated by workflows.

By implementing SciPipe as a library in an existing language, the language’s ecosystem of tooling, editor support and third-party libraries can be directly used to avoid “reinventing the wheel” in these areas. By leveraging the built-in concurrency features of Go, such as go-routines and channels, the developed code base has been kept small compared with similar tools, and also does not require external dependencies for basic usage (some external tools are used for optional features like PDF generation and graph plotting). This means that the code base should be possible to maintain for a single developer or small team, and that the code base is practical to include in workflow developers’ own source code repositories, in order to future-proof the functionality of workflows.

Below, we first briefly describe how SciPipe workflows are created. We then describe in some detail the features of SciPipe that are the most novel or improves most upon existing tools, followed by a few more commonplace technical considerations. We finally demonstrate the usefulness of SciPipe by applying it to a set of case study workflows in machine learning for drug discovery and next-generation sequencing genomics and transcriptomics.

### Writing workflows with SciPipe

SciPipe workflows are written as Go programs, in files ending with the .go extension. As such, they require the Go tool chain to be installed for compiling and running them. The Go programs can be either compiled to self-contained executable files with the `go build` command, or run directly, using the `go run` command.

The simplest way to write a SciPipe program is to write the workflow definition in the program’s `main()` function, which is executed when running the compiled executable file, or running the file as script with `go run`. An example workflow written in this way is shown in in figure 1, which provides a simple example workflow consisting of three processes, demonstrating a few of the basic features of SciPipe. The first process writes a string of DNA to a file, the second computes the base complement, and the last process reverses the string. All in all, the workflow computes the reverse base complement of the initial string.

**Figure 1:**
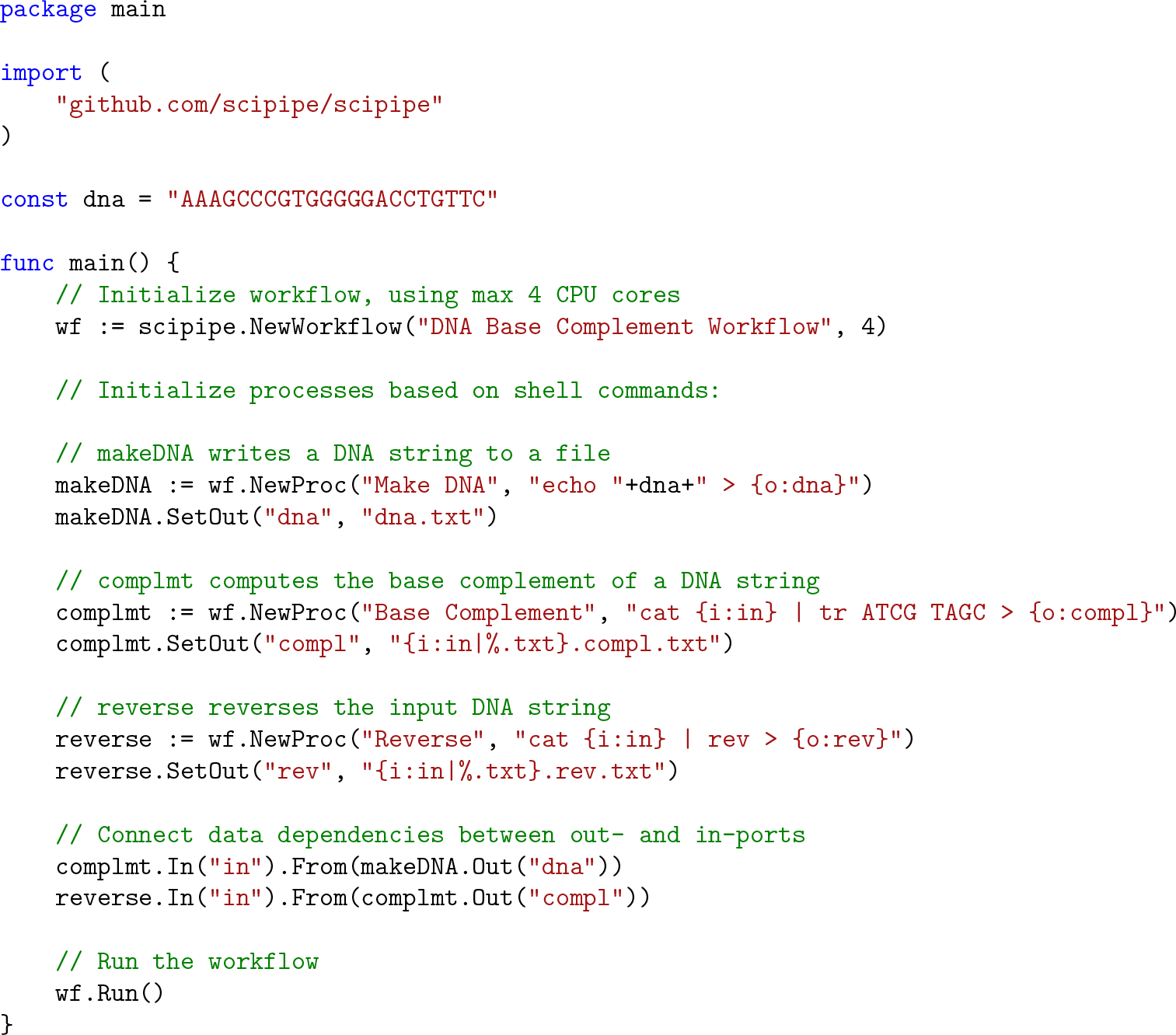
A simple example workflow implemented with SciPipe. The workflow computes the reverse base complement of a string of DNA, using standard UNIX tools. The workflow is a Go program and is supposed to be saved in a file with the .go extension and executed with the go run command. On line 4, the SciPipe library is imported, to be later accessed as `scipipe`. On line 7, a short string of DNA is defined. On line 9-33, the full workflow is implemented in the program’s main() function, meaning that it will be executed when the resulting program is executed. On line 11, a new workflow object (or “struct” in Go terms) is initiated with a name and the maximum number of cores to use. On lines 15-25, the workflow components, or *processes*, are initiated, each with a name and a shell command pattern. Input file names are defined with a placeholder on the form {`i:INPORTNAME`} and outputs on the form {`o:OUTPORTNAME`}. The port-name will be used later to access the corresponding ports for setting up data dependencies. On line 16, a component that writes the previously defined DNA string to a file is initiated, and on line 17, the file path pattern for the out-port *dna* is defined (in this case a static file name). On line 20, a component that translates each DNA base to its complementary counterpart is initiated. On line 21, the file path pattern for its only out-port is defined. In this case, reusing the file path of the file it will receive on its in-port named *in*, thus the {i:in} part. The %.txt part removes .txt from the input path. On line 24, a component that will reverse the DNA string is initiated. On lines 27-29, data dependencies are defined via the in-and out-ports defined earlier as part of the shell command patterns. On line 32, the workflow is being run.

As can be seen in figure 1 on line 11, a workflow object (or struct, in Go terminology) is first initialized, with a name and a setting for the maximum number of tasks to run at a time. Furthermore, on line 15-19, processes are defined with the `Workflow.NewProc()` method on the workflow struct, with name and a command pattern which is very similar to the Bash shell command that would be used to run a command manually, but where concrete file names have been replaced with placeholders, on the form {`i:INPORTNAME`}, {`o:OUTPORTNAME`} or {`p:PARAMETERNAME`}. These placeholders define input and output files, as well as parameter values, and works as a sort of templates, that will be replaced with concrete values as concrete tasks are scheduled and executed.

As can be seen on lines 17, 21 and 25, output paths to use for output files are defined using the `Process.SetOut()` method, taking an out-port name and a pattern for how to generate the path. For simple workflows this can be just a static file name, but for more complex workflows with processes that produce more than one output on the same port - e.g. by processing different input files, or using different sets of parameters - it is often best to reuse some of the input paths and parameter values configured earlier in the command pattern to generate a unique path for each output.

Finally, on lines 27-29, we see how in-ports and out-ports are connected in order to define the data depen-dencies between tasks. Here, the in-port and out-port names used in the placeholders in the command pattern described above, are used to access the corresponding in-ports and out-ports, and making connections between them, with a syntax on the general form of `InPort.From(OutPort)`.

The last thing needed to do to run the workflow, is seen on line 32, where the `Workflow.Run()` method is executed. Provided that the workflow code in figure 1 is saved in a file named `workflow.go`, it can be run using the go run command, like so:

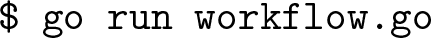

This will then produce three output files and one accompanying audit log for each file, which we can be seen by listing the files in a terminal:

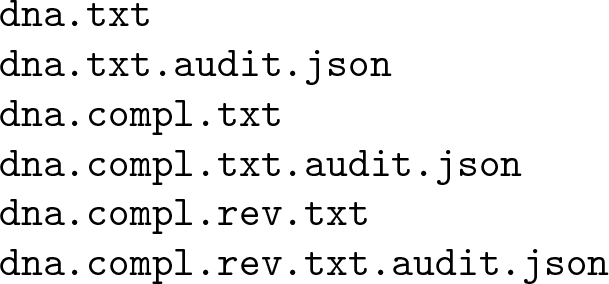

The file dna.txtshould now contain the string AAAGCCCGTGGGGGACCTGTTC, and `dna.compl.rev.txt` should contain GAACAGGTCCCCCACGGGCTTT, which is the reverse base complement of the first string. In the last file above, the full audit log for this minimal workflow can be found. An example content of this file is shown in figure 2.

**Figure 2:**
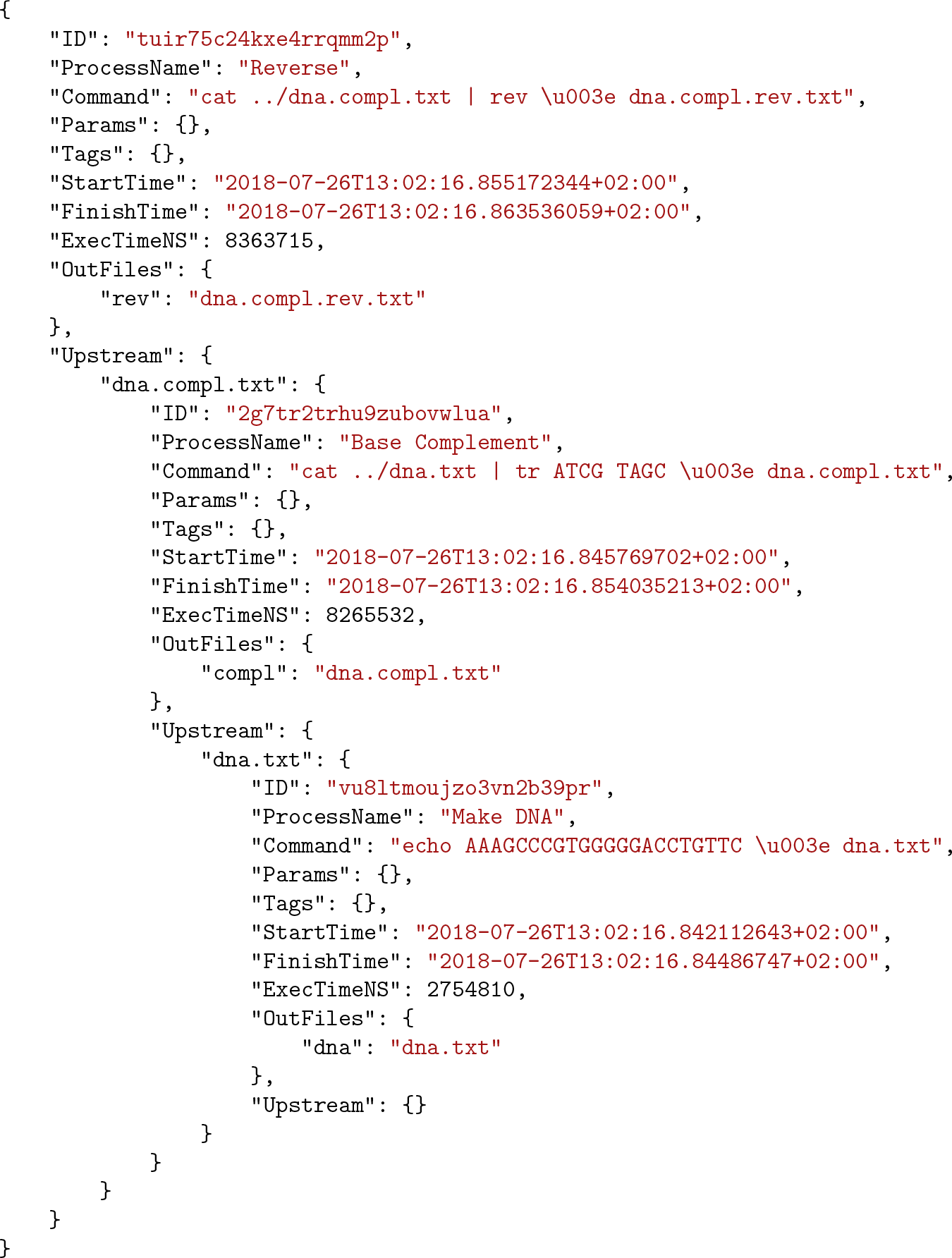
Example audit log file in JSON format [23], for a file produced by a SciPipe workflow. The workflow used to produce this audit log in particular, is the one in figure 1. The audit information is hierarchical, with each level representing a step in the workflow. The first level contains meta-data about the task executed last, to produce the output file that this audit log refers to. The field `Upstream` on each level, contains a list of all upstream task of the current task, indexed by the file paths that each of the upstream tasks did produce, and which was subsequently used by the current task. Each task is given a globally unique ID, which helps to deduplicate any duplicate occurrences of tasks, when converting the log to other representations. Execution time is given in nanoseconds. Note that input paths in the command field, is prepended with../, compared to how they appear in the Upstream field. This is because each task is executed in a temporary directory created directly under the workflow’s main execution directory, meaning that to access existing data files, it has to first navigate up one step out of this temporary directory.

In this code example, it can be seen that both of the commands we executed are available, and also that the *Reverse* process lists its “upstream” processes, which are indexed by the input file names in its command. Thus, under the dna.compl.txt input file, we find the *Base Complement* process together with its meta-data, and one further upstream process (the *Make DNA* process). This hierarchic structure of the audit log ensures that the complete audit trace, including all commands contributing to the production of an output file, is available for each output file from the workflow.

More information about how to write workflows with SciPipe is available on the documentation website [24]. Note that the full documentation on this website is also available in a folder named docs inside the SciPipe Git repository, which ensures that documentation for the version currently used, is always available.

### Dynamic scheduling

Since SciPipe is built on the principles from Flow-based programming (see the methods section for more details), a SciPipe program consists of independently and concurrently running processes, which schedule new tasks continually during the workflow run. This is here referred to as *dynamic scheduling*. This means that it is possible to create a process that obtains a value and passes it on to a downstream process as a parameter, so that new tasks can be scheduled with it. This feature is important in machine learning workflows, where hyper parameter tuning is often employed to find an optimal value of a parameter, such as cost for Support Vector Machines (SVM), which is then used to parametrize the final training part of the workflow.

### Reusable components

Based on principles from Flow-based programming, the workflow graph in SciPipe is defined by making connections between port objects bound to processes. This enables to keep the dependency graph definition *separate* from the process definitions. This is in contrast to other ways of connecting dataflow processes, such as with dataflow variables, which are shared between process definitions. This makes processes in flow-based programming fully self-contained, meaning that libraries of reusable components can be created and that components can be loaded on-demand when creating new workflows. A graphical comparison between dependencies defined with dataflow variables and flow-based programming ports, is shown in figure 3.

**Figure 3:**
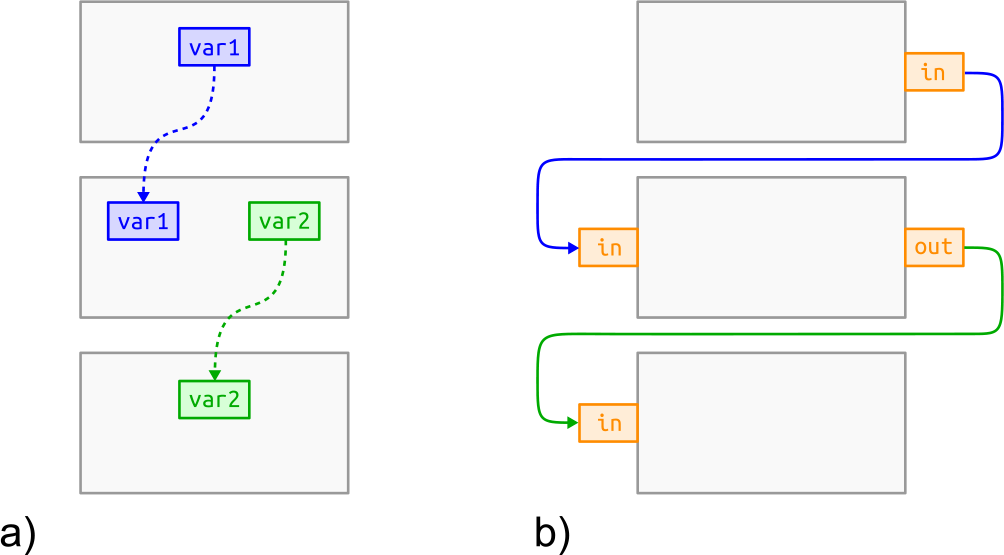
Comparison between dataflow variables and Flow-based programming ports in terms of dependency definition. a) shows how dataflow variables (blue and green) shared between processes (in gray) make the processes tightly coupled. In other words, process-and network definitions get intermixed. b) shows how ports (in orange) bound to processes in Flow-based programming allows keeping the network definition separate from process definitions. This enables processes to be reconnected freely without changing their internals.

### Running subsets of workflows

With pull-based workflow tools like Snakemake or Luigi, it is easy to on-demand reproduce a particular output file, since the scheduling mechanism is optimized for the use case of asking for a specific file and calculating all the tasks required to be executed based on that.

With push-based workflow tools though, reproducing a specific set of files without running the full workflow is not always straight-forward. This is a natural consequence of the push-based scheduling strategy, and dataflow in particular, as the identities and quantities of output files might not be known before the workflow is run.

SciPipe provides a mechanism for partly solving this lack of “on demand file generation” in push-based dataflow tools, by allowing to reproduce all files of a specified process, on-demand. That is, the user can tell the workflow to run all processes in the workflow upstream of, and including, a specified process, while skipping processes downstream of it.

This has turned out very useful when iteratively refactoring or developing new pipelines. When a part in the middle of a long sequence of processes need to be changed, it is helpful to be able to test-run the workflow up to that particular process only, not the whole workflow, to speed up the development iteration cycle.

### Other characteristics

Below are a few technical characteristics and considerations that are not necessarily unique to SciPipe, but could be of interest to potential users assessing whether SciPipe fits their use cases.

### Data centric audit log

The audit log feature in SciPipe collects meta data about every executed task (concrete shell command invocation) which is passed along with every file that is processed in the workflow. It writes a file in the ubiquitous JSON format, with the full trace of tasks executed for every output in the workflow, with the same name as the output file in question but with the additional file extension `.audit.json.` Thus, for every output in the workflow, it is possible to check the full record of shell commands used to produce it. An example audit log file can be seen in figure 2.

This data-oriented provenance reporting contrasts to provenance reports common in many workflow tools, which often provide one report per workflow run only, meaning that the link between data and provenance report is not as direct.

The audit log feature in SciPipe in many aspects reflects the recommendations in [25] for developing provenance reporting in workflows, such as producing a coherent, accurate, inspectable record for every output data item from the workflow. By producing provenance records for each data output rather than for the full workflow only, SciPipe could provide a basis for the problem of iteratively developing workflow variants, as outlined in [26].

SciPipe also loads any existing provenance reports for existing files that it uses, and merges these with the provenance information from its own execution. This means that even if a chain of processing is spread over multiple SciPipe workflow scripts, and executed at different times by different users, the full provenance record is still being kept and assembled, as long as all workflow steps were executed using SciPipe shell command processes. The main limitation to this “follow the data” approach, is for data generated externally to the workflow, or by SciPipe components implemented in Go. For external processes, it is up to the external process to generate any reporting. For Go-based components in SciPipe, these can not currently dump a textual version of the Go code executed. This constitutes an area of future development.

SciPipe provides experimental support for converting the JSON-structure into reports in HTML and TeX format, or into executable Bash scripts that can reproduce the file which the audit report describes from available inputs or from scratch. These tools are available in the `scipipe` helper command. The `TeX` report can be easily further converted to PDF using the `pdflatex` command of the `pdfTex` software [27]. An example of such a PDF report, is shown in figure 4, which was generated from the audit report for the last file generated by the code example in figure 1.

**Figure 4:**
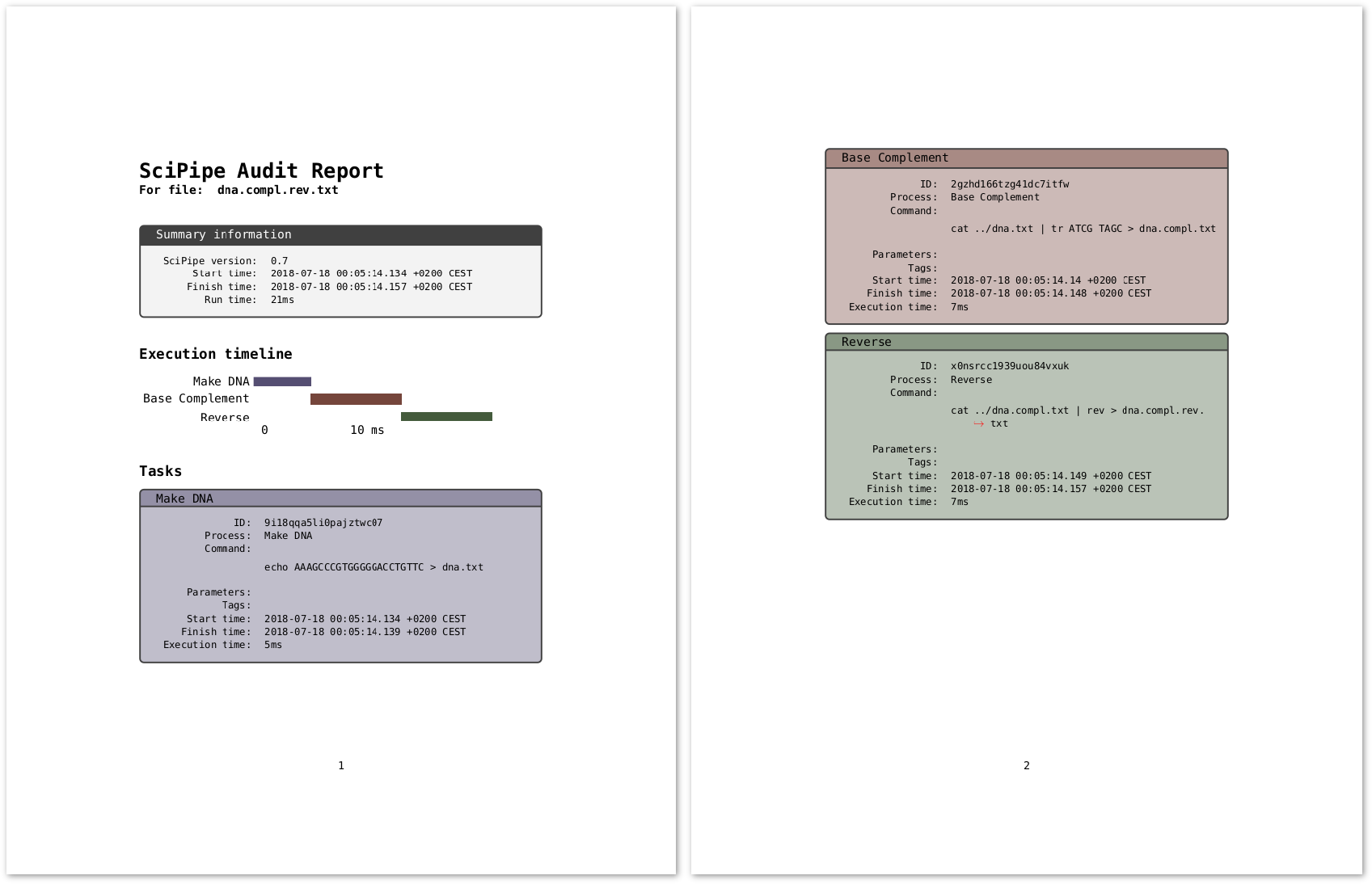
Audit report for the last file generated by the code example in figure 1, converted to TeX with SciPipe’s experimental `audit2tex` feature and then converted to PDF with pdfTeX. In the top, the PDF file includes summary information about the SciPipe version used and the total execution time. After this follows an execution time line, in a gantt-chart style, that shows the relative execution times of individual tasks in a graphical way. After this follows a comprehensive list of tables with information for each task executed towards producing the file for which the audit report belongs. The task boxes are color coded and ordered in the same way that the tasks appear in the timeline.

### Atomic writes

SciPipe ensures that cancelled workflow runs do not result in half-written output files being mistaken for finished ones. It does this by executing each task in a temporary folder, and moving all newly created files into their final location *after* the task is finished. By using a folder for the execution, any extra files created by a tool that are not explicitly configured by the workflow system, are captured and treated in an atomic way. Examples of where this is needed is for the five extra files created by `bwa index` [28], when indexing a reference genome in FASTA format.

### Streaming support

In data intensive fields like Next-Generation Sequencing, it is common that intermediate steps of pipelines produce large amounts of intermediate data, often multiplying the storage requirements considerably compared to the raw data from sequencing machines [29]. To help ease these storage requirements, SciPipe provides the ability to optionally stream data between two tasks via Random Access Memory (RAM) instead of saving to disk between task executions. This approach has two benefits. Firstly, the data does not need to be stored on disk, which can lessen the storage requirements considerably. Secondly, it enables the downstream task to start processing the data from the upstream task immediately as soon as the first task has started to produce partial output. It thus enables to achieve *pipeline parallelism* in addition to *data parallelism*, and can thereby shorten the total execution time of the pipeline.

### Flexible file naming and data “caching”

SciPipe allows flexible naming of the file path of every intermediate output in the workflow, based on input file names and parameter values passed to a process. This enables creating meaningful file naming schemes, to make it easy to manually explore and sanity-check outputs from workflows.

Configuring a custom file naming scheme is not required though. If no file path is configured, SciPipe will automatically create a file path that ensures that two tasks with different parameters or input data will never clash, and that two tasks with the same command signature, parameters and input-files, will reuse the same cached data.

### Known limitations

Below we list some design decisions and known limitations of SciPipe that might affect the decision whether to use SciPipe for a particular use case or not.

Firstly, the fact that writing SciPipe workflows requires some basic knowledge of the Go programming language, can be off-putting to users who are not well acquainted with programming. Go code, although having taken inspiration from scripting languages, is still markedly more verbose and low-level in nature than Python, and can take a little longer to get used to.

Secondly, the level of integration with HPC resource managers is currently quite basic compared to some other workflow tools. The SLURM resource manager can readily be used by using the `Prepend` field on processes to add a string with a call to the `salloc` SLURM command, but more advanced HPC integration is planned to be addressed in upcoming versions.

Furthermore, reproducing specific output files is not as natural and easy as with pull-based tools like Snake-make, although SciPipe provides a mechanism to partly resolve this problem.

Finally, SciPipe does not yet support integration with the Common Workflow Language [30], for interoperability of workflows. This is a prioritized area for future development.

## Case Studies

To demonstrate the usefulness of SciPipe, we have used it to implement a number of representative pipelines from drug discovery and bioinformatics with different characteristics and hence requirements on the workflow system. These workflows are available in a dedicated git repository on GitHub [31].

### Machine learning pipeline in drug discovery

The initial motivation for building SciPipe stemmed from problems encountered with complex dynamic workflows in machine learning for drug discovery applications. It was thus quite natural to implement an example of such a workflow in SciPipe. To this end we re-implemented a workflow implemented previously for the SciLuigi library [20], which was itself based on an earlier study [32].

In short, this workflow trains predictive models using the LIBLINEAR software [33] with molecules represented by the signature descriptor [34]. For linear SVM a cost parameter needs to be tuned, and we tested 15 values (0.0001, 0.0005, 0.001, 0.005, 0.01, 0.05, 0.1, 0.25, 0.5, 0.75, 1, 2, 3, 4, 5) in a 10-fold cross-validated parameter sweep. Five different training set sizes (500, 1000, 2000, 4000, 8000) were tested and evaluated with a test set size of 1000. The raw data set consists of 10,000 logarithmic solubility values chosen randomly from a dataset extracted from PubChem [35] according to details in [20]. The workflow is schematically shown in figure 5 and was plotted using SciPipe’s built-in plotting function. The figure has been modified for clarity by collapsing the individual branches of the parameter sweeps and cross validation folds, as well as by manually making the layout more compact.

The implementation in SciPipe was done by creating components which are defined in separate files (named `comp_COMPONENTNAME` in the repository), which can thus be reused in other workflows. This shows how SciPipe can successfully be used to create workflows based on reusable, externally defined components.

**Figure 5:**
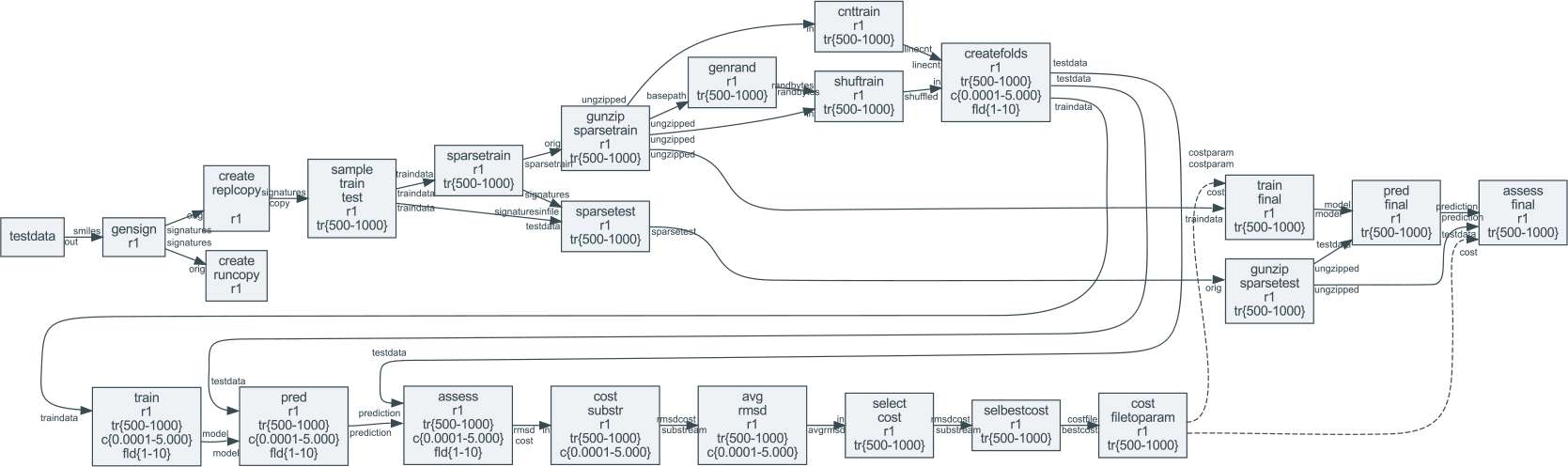
Directed graph of the machine learning drug discovery case study workflow, plotted with SciPipe’s workflow plotting function. The graph has been modified for clarity by collapsing the individual branches of the parameter sweeps and cross validation fold-generation. The layout has also been manually made more compact to be viewable in print. The collapsed branches are indicated by intervals in the box labels. `tr {500-8000}` represent branching into training dataset sizes `500, 1000, 2000, 4000, 8000. c {0.0001-5.0000}` represent cost values `0.0001, 0.0005, 0.001, 0.005, 0.01, 0.05, 0.1, 0.25, 0.5, 0.75, 1, 2, 3, 4` and `5`, while `fld 1-10` represent cross validation folds 1-10. Nodes represent processes, while edges represent data dependencies. The labels on the edge heads and tails represent ports. Solid lines represent data dependencies via files, while dashed lines represent data dependencies via parameters, which are not persisted to file, only transmitted via RAM.

The fact that SciPipe supports parametrization of workflow steps with values obtained during the workflow run, meant that the full workflow could be kept in a single workflow definition, in one file. This also made it possible to create audit logs for the full workflow execution for the final output files, and to create the automatically plotted workflow graph shown in figure 5. This is in contrast to the SciLuigi implementation, where the parameter sweep to find the optimal cost, and the final training, had to be implemented in separate workflow files (`wffindcost.py` and `wfmm.py` in [36]), and executed as a large number of completely separate workflow runs (one for each dataset size) which meant that logging became fragmented into a large number of disparate log files.

### Genomics cancer-analysis pipeline

Sarek [37] is an open-source analysis pipeline to detect germ-line or somatic variants from whole genome sequencing, developed by the National Genomics Infrastructure and National Bioinformatics Infrastructure Sweden which are both platforms at Science for Life Laboratory.

To test and demonstrate the applicability of SciPipe to genomics use cases the pre-processing part of the Sarek pipeline was implemented in SciPipe. See figure 6 for a directed process graph of the workflow, plotted with SciPipe’s workflow plotting function.

**Figure 6:**
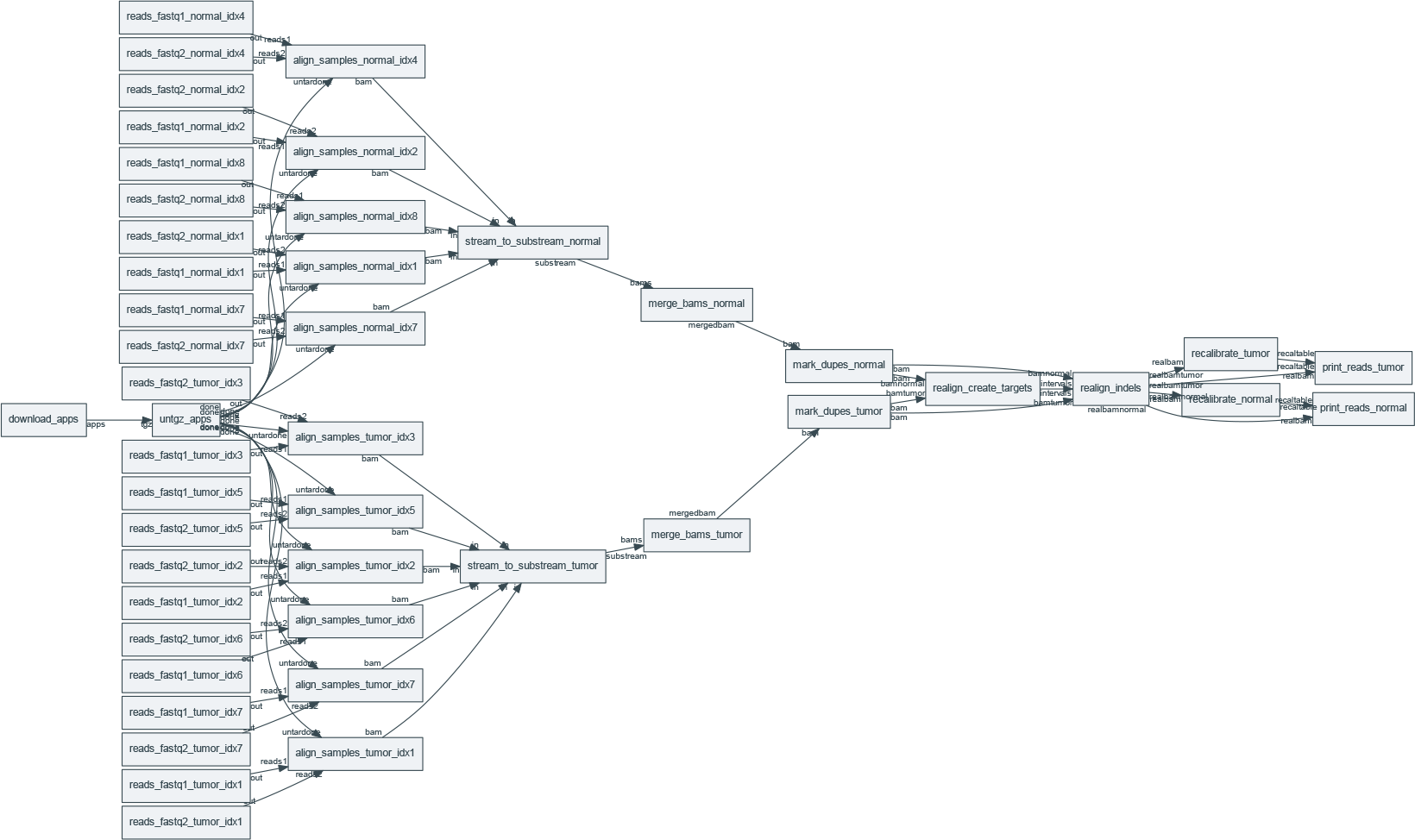
Directed graph of workflow processes in the Genomics / Cancer Analysis pre-processing pipeline, plotted with SciPipe’s workflow plotting function. Nodes represent processes, while edges represent data de-pendencies. The labels on the edge heads and tails represent ports.

The test data in the test workflow consists of multiple samples of normal and tumor pairs. The workflow starts with aligning each sample to a reference genome using BWA [28] and forwarding the results to Samtools [38] which saves the result as a sorted BAM file. After each sample has been aligned, Samtools is again used, to merge the normal-and tumor samples into a one BAM [38] file for tumor samples, and one for normal. Picard [39] is then used to mark duplicate reads in both the normal-and tumor sample BAM files, whereafter GATK [40] is used to recalibrate the quality scores of all reads. The outcome of the workflow is two BAM files; one containing all the normal samples and one containing all the tumor samples.

Genomics tools and pipelines have their own set of requirements, which was shown by the fact that some aspects of SciPipe had to be modified in order to ease development of this pipeline. In particular, many genomics tools produce additional output files apart from those specified on the command-line. One example of this is the extra files produced by BWA when indexing a reference genome in FASTA format. The `bwa index` command produces some five files, which are not explicitly defined on the command-line (with the extensions of `.bwt, .pac, .ann, .amb and .sa`). Based on this realization, SciPipe was amended with a folder-based execution mechanism which executes each task in a temporary folder, that keeps all output files separate from the main output directory until the whole task has completed. This ensures that also files that are not explicitly defined and handled by SciPipe, are also captured and handled in an atomic manner, so that finished and unfinished output files are always properly separated.

Furthermore, agile development of genomic tools often requires being able to see the full command that is used to execute a tool, because of the many options that are available to many bioinformatics tools. This workflow was thus implemented with ad-hoc commands, which are defined in-line in the workflow. The ability to do this shows that SciPipe supports different ways of defining components, depending on what fits the use case best.

The successful implementation of this genomics pipeline in SciPipe, thus both ensures and shows that SciPipe is works well for tools common in genomics.

### RNA-seq / transcriptomics pipeline

**Figure 7:**
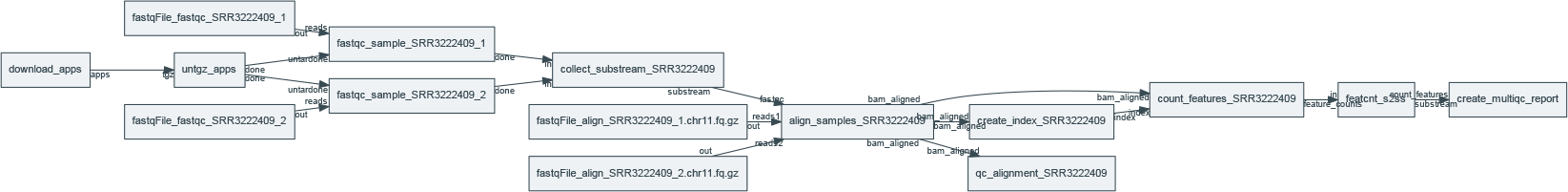
Directed graph of workflow processes in the RNA-Seq Pre-processing workflow, plotted with SciPipe’s workflow plotting function. Nodes represent processes, while edges represent data dependencies. The labels on the edge heads and tails represent ports.

To test the ability of SciPipe to work with software used in transcriptomics, some of the initial steps of a generic RNA-sequencing workflow were also implemented in SciPipe. Common steps that are needed in transcriptomics is to run quality controls and generate reports of the analysis steps.

The RNA-seq case study pipeline implemented for this paper uses FastQC [41] to evaluate the quality of the raw data being used in the analysis before aligning the data using STAR [42]. After the alignment is done it is evaluated using QualiMap [43], while the Subread package [44] is used to do a feature counting.

The final step of the workflow is to combine all the previous steps for a composite analysis using MultiQC [45], which will summarize the quality of both the raw data and the result of the alignment into a single quality report. See figure 7 for a directed process graph of the workflow, plotted with SciPipe’s workflow plotting function.

The successful implementation of this transcriptomics workflow in SciPipe ensures that SciPipe works well for different types of bioinformatics workflows and is not limited to one specific sub-field of bioinformatics.

## Conclusions

SciPipe is a programming library that provides a way to write complex and dynamic pipelines in bioinformatics, cheminformatics, and more generally in data science and machine learning pipelines involving command-line applications.

Dynamic scheduling allows parametrizing new tasks with values obtained during the workflow run, and the Flow-based programming principles of separate network definition and named ports allow creating a library of reusable components. By having access to the full power of the Go programming language to define workflows, existing tooling is leveraged.

Scipipe adopts state-of-the art strategies for achieving atomic writes, caching of intermediate files and a data-centric audit log feature that allows identifying the full execution trace for each output, that can be exported into either human-readable HTML or TeX/PDF formats, or executable Bash-scripts.

SciPipe also provides some features not commonly found in many tools such as support for streaming via Unix named pipes, ability to run push-based workflows up to a specific stage of the workflow, and flexible support for file naming of intermediate data files generated by workflows. SciPipe workflows can also be compiled into standalone executables, making deployment of pipelines maximally easy, requiring only Bash and any external command-line tools used, to be present on the target machine.

By being a small library without required external dependencies apart from the Go tool chain and Bash, SciPipe is expected to be possible to be maintained and developed in the future even without a large team or organization backing it.

The applicability of SciPipe for cheminformatics, genomics and transcriptomics pipelines has been demonstrated with case study workflows in these fields.

## Methods

### The Go Programming Language

The Go Programming Language (referred to as just “Go”) was developed by Robert Griesemer, Rob Pike and Ken Thompson at Google, to provide a statically typed and compiled language that makes it easier to build highly concurrent programs, that can also make good use of multiple CPU cores (i.e. “parallel program”), than what is the case in widespread compiled languages like C++ [46]. It tries to provide this by providing a small, simple language, with concurrency primitives — go-routines and channels — built-in to the language. Go-routines, which are so called light-weight threads, are automatically mapped, or multiplexed, onto physical threads in the operating system. This means that very large numbers of go-routines can be created while maintaining a low number of operating system threads, such as one per CPU core on the computer at hand. This makes Go an ideal choice for problems where many asynchronously running processes need to be handled at the same time, or “concurrently”, and for making efficient use of multi-core CPUs.

The Go compiler is statically linking all its code as part of the compilation. This means that all dependent libraries are compiled into the executable file. Because of this, SciPipe workflows can be compiled into self-contained executable files without external dependencies apart from the Bash shell and any external command line tools used by the workflow. This makes deploying Go programs (and SciPipe workflows) to production very easy.

Go programs are very performant, often an order of magnitude faster than interpreted languages like Python, and in the same order of magnitude as the fastest languages, like C, C++ and Java [47].

### Dataflow and Flow-based programming

Dataflow is a programming paradigm oriented around the idea of independent, asynchronously running processes, that only talk to each other by passing data between each other. This data passing can happen in different ways, such as via dataflow variables, or via first-in-first-out channels.

Flow-Based Programming (FBP) [48] is a paradigm for programming developed by John Paul Morrison at IBM in the late 60s / early 70s, to provide a composable way to assemble programs to be run at mainframe computers at customers such as large banks.

It is a specialized version of dataflow, adding the ideas of separate network definition, named ports, channels with bounded buffers and information packets (representing the data) with defined lifetimes. Just as in dataflow, the idea is to divide a program into independent processing units called “processes”, which are allowed to communicate with the outside world and other processes solely via message passing. In FBP, this is always done over channels with bounded buffers which are connected to named ports on the processes. Importantly, the network of processes and channels is in FBP described “separate” from the process implementations, meaning that the network of processes and channels can be reconnected freely without changing the internals of processes.

This strict separation of the processes, the separation of network structure from processing units, and the loosely-coupled nature of its only way of communication with the outside world (message passing over channels) makes flow-based programs extremely composable, and naturally component-oriented. Any process can always be replaced with any other process that supports the same format of the information packets on its in-ports and out-ports.

Furthermore, since the processes run asynchronously, FBP is, just like Go, very well suited to make efficient use of multi-core CPUs, where each processing unit can suitably be placed in its own thread or co-routine to spread out on the available CPU-cores on the computer. FBP has a natural connection to workflow systems, where the computing network in an FBP program can be likened to the network of dependencies between data and processing components in a workflow [20]. SciPipe leverages the principles of separate network definition and named ports on processes. SciPipe has also taken some inspiration for its API design from the GoFlow [49] Go-based flow-based programming framework.

## Availability of supporting source code and requirements

- Project name: SciPipe
- Documentation and project home page: http://scipipe.org
- Source code repository: https://github.com/scipipe/scipipe
- Persistent source code archive: https://doi.org/10.5281/zenodo.1157941
- Case study workflows: https://github.com/pharmbio/scipipe-demo
- Operating system(s): Linux, Unix, Mac
- Other requirements: Bash, GraphViz (for workflow graph plotting), LaTeX (for PDF generation)
- License: MIT

## Availability of supporting data

- The raw data for the machine learning cheminformatics demonstration pipeline is available at: https://doi.org/10.5281/zenodo.1324443
- The applications for the machine learning in drug discovery case study is available at: https://doi.org/10.6084/m9.figshare.3985674.v1
- The raw data and tools for the genomics and transcriptomics workflows are available at: https://doi.org/10.5281/zenodo.1324426

## Declarations

### List of abbreviations

- API: Application programming interface
- CPU: Central processing unit (the core part of every computer)
- DSL: Domain-specific language
- FBP: Flow-based programming
- HPC: High-performance computing
- RAM: Random access memory
- SVM: Support vector machine

### Ethics approval and consent to participate

Not applicable.

### Consent for publication

Not applicable.

## Authors’ Contributions

OS and SL conceived the project and the idea of component-based workflow design. SL came up with the idea of using Go and Flow-based programming principles to implement a workflow library, designed and implemented the SciPipe library, and implemented case study workflows. MD contributed to the API design and implemented case study workflows. JA implemented the TeX/PDF reporting function and contributed to case study workflows. OS supervised the project. All authors read and approved the manuscript.

## Acknowledgments

We thank Egon Elbre, Johan Dahlberg and Johan Viklund for valuable feedback and suggestions regarding the workflow API. We thank Rolf Lampa for valuable suggestions on the audit log feature. We also thank colleagues at pharmb.io and on the SciLifeLab slack for helpful feedback on the API, and users on the Flow-based programming mailing list for encouraging feedback.

## Bibliography

1 Nils Gehlenborg, Seán I O’donoghue, Nitin S Baliga, Alexander Goesmann, Matthew A Hibbs, Hiroaki Kitano, Oliver Kohlbacher, Heiko Neuweger, Reinhard Schneider, Dan Tenenbaum, et al. Visualization of omics data for systems biology. Nature methods, 7(3s):S56, 2010.

2 Marylyn D Ritchie, Emily R Holzinger, Ruowang Li, Sarah A Pendergrass, and Dokyoon Kim. Methods of integrating data to uncover genotype–phenotype interactions. Nature Reviews Genetics, 16(2):85, 2015.

3 Vivien Marx. Biology: The big challenges of big data. Nature, 498(7453):255–260, 2013.

4 Zachary D. Stephens, Skylar Y. Lee, Faraz Faghri, Roy H. Campbell, Chengxiang Zhai, Miles J. Efron, Ravishankar Iyer, Michael C. Schatz, Saurabh Sinha, and Gene E. Robinson. Big data: Astronomical or genomical? PLoS Biology, 13(7):1–11, 2015.

5 O. Spjuth, E. Bongcam-Rudloff, G.C. Hern?ndez, L. Forer, M. Giovacchini, R.V. Guimera, A. Kallio, E. Korpelainen, M.M. Ka?dula, M. Krachunov, D.P. Kreil, O. Kulev, P.P. ?abaj, S. Lampa, L. Pireddu, S. Sch?nherr, A. Siretskiy, and D. Vassilev. Experiences with workflows for automating data-intensive bioinformatics. Biology Direct, 10(1), 2015.

6 Daniel Blankenberg, Gregory Von Kuster, Nathaniel Coraor, Guruprasad Ananda, Ross Lazarus, Mary Mangan, Anton Nekrutenko, and James Taylor. Galaxy: A Web-Based Genome Analysis Tool for Experimentalists. John Wiley & Sons, Inc., Hoboken, 2010.

7 Belinda Giardine, Cathy Riemer, Ross C. Hardison, Richard Burhans, Laura Elnitski, Prachi Shah, Yi Zhang, Daniel Blankenberg, Istvan Albert, James Taylor, Webb Miller, W. James Kent, and Anton Nekrutenko. Galaxy: A platform for interactive large-scale genome analysis. Genome Res., 15(10):1451–1455, 2005.

8 B. Giardine, C. Riemer, R. C. Hardison, R. Burhans, L. Elnitski, P. Shah, Y. Zhang, D. Blankenberg, I. Albert, J. Taylor, W. Miller, W. J. Kent, and A. Nekrutenko. Galaxy: a platform for interactive large-scale genome analysis. Genome Res., 15, 2005.

9 Adam A. Hunter, Andrew B. Macgregor, Tamas O. Szabo, Crispin A. Wellington, and Matthew I. Bellgard. Yabi: An online research environment for grid, high performance and cloud computing. Source Code Biol. Med., 7(1):1–10, 2012.

10 Johannes Köster and Sven Rahmann. Snakemake—a scalable bioinformatics workflow engine. Bioinfor-matics, 28(19):2520–2522, 2012.

11 Paolo Di Tommaso, Maria Chatzou, Evan W Floden, Pablo Prieto Barja, Emilio Palumbo, and Cedric Notredame. Nextflow enables reproducible computational workflows. Nature Biotech, 35(4):316–319, 2017.

12 Simon P. Sadedin, Bernard Pope, and Alicia Oshlack. Bpipe: a tool for running and managing bioinformatics pipelines. Bioinformatics, 28(11):1525–1526, 2012.

13 Jörgen Brandt, Marc Bux, and Ulf Leser. Cuneiform: a functional language for large scale scientific data analysis. In EDBT/ICDT Workshops, pages 7–16, 2015.

14 Jon Ander Novella, Payam Emami Khoonsari, Stephanie Herman, Daniel Whitenack, Marco Capuccini, Joachim Burman, Kim Kultima, and Ola Spjuth. Container-based bioinformatics with pachyderm. bioRxiv, 2018.

15 Erik Bernhardsson, Elias Freider, and Arash Rouhani. spotify/luigi-GitHub. https://github.com/spotify/luigi. [Online; Accessed 3-July-2018].

16 Yolanda Gil and Varun Ratnakar. Dynamically generated metadata and replanning by interleaving workflow generation and execution. In Semantic Computing (ICSC), 2016 IEEE Tenth International Conference on, pages 272–276. IEEE, 2016.

17 David K Rensin. Kubernetes-scheduling the future at cloud scale. 2015.

18 Johan Dahlberg, Johan Hermansson, Steinar Sturlaugsson, and Pontus Larsson. Arteria: An automation system for a sequencing core facility. bioRxiv, 2017.

19 Jörgen Brandt, Wolfgang Reisig, and Ulf Leser. Computation semantics of the functional scientific workflow language cuneiform. Journal of Functional Programming, 27:e22, 2017.

20 Samuel Lampa, Jonathan Alvarsson, and Ola Spjuth. Towards agile large-scale predictive modelling in drug discovery with flow-based programming design principles. Journal of Cheminformatics, 8(1):67, 2016.

21 Samuel Lampa. SciPipe source code repository at GitHub. https://github.com/scipipe/scipipe. [Online; Accessed 4-July-2018].

22 Samuel Lampa, Martin Czugan, and Jonathan Alvarsson. scipipe/scipipe latest version - zenodo. https://doi.org/10.5281/zenodo.1157941, July 2018.

23 Douglas Crockford. JSON website. http://json.org/. [Online; Accessed 16-July-2018].

24 Samuel Lampa. SciPipe documentation. http://scipipe.org. [Online; Accessed 5-July-2018].

25 Yolanda Gil and Daniel Garijo. Towards Automating Data Narratives. Proceedings of the 22nd International Conference on Intelligent User Interfaces - IUI’17, (February):565–576, 2017.

26 Lucas A M C Carvalho, Bakinam T Essawy, Daniel Garijo, Claudia Bauzer Medeiros, and Yolanda Gil. Requirements for Supporting the Iterative Exploration of Scientific Workflow Variants. 2017 Workshop on Capturing Scientiftc Knowledge (SciKnow), 2017.

27 Peter Breitenlohner and Han The Thanh. pdfTeX. http://www.tug.org/applications/pdftex. [Online; Accessed 25-July-2018].

28 Heng Li and Richard Durbin. Fast and accurate short read alignment with Burrows-Wheeler transform. Bioinformatics, 25(14):1754–1760, 15 July 2009.

29 Martin Dahlö, Douglas G Scofield, Wesley Schaal, and Ola Spjuth. Tracking the ngs revolution: managing life science research on shared high-performance computing clusters. GigaScience, 7(5):giy028, 2018.

30 Peter Amstutz, Michael R. Crusoe, Nebojša Tijanić, Brad Chapman, John Chilton, Michael Heuer, Andrey Kartashov, Dan Leehr, Hervé Ménager, Maya Nedeljkovich, Matt Scales, Stian Soiland-Reyes, and Luka Stojanovic. Common Workflow Language, v1.0. 7 2016.

31 Samuel Lampa, Martin Dahlö, Jonathan Alvarsson, and Ola Spjuth. SciPipe Demonstration workflows source code repository at GitHub. https://github.com/pharmbio/scipipe-demo. [Online; Accessed 26-July-2018].

32 Jonathan Alvarsson, Samuel Lampa, Wesley Schaal, Claes Andersson, Jarl E S Wikberg, and Ola Spjuth. Large-scale ligand-based predictive modelling using support vector machines. Journal of Cheminformatics, 8, 2016.

33 Rong-En Fan, Kai-Wei Chang, Cho-Jui Hsieh, Xiang-Rui Wang, and Chih-Jen Lin. LIBLINEAR: A Library for Large Linear Classification. Journal of Machine Learning Research, 9(2008):1871–1874, 2008.

34 Jean-Loup Faulon, Donald P Visco, and Ramdas S Pophale. The signature molecular descriptor. 1. using extended valence sequences in qsar and qspr studies. Journal of chemical information and computer sciences, 43(3):707–720, 2003.

35 National Center for Biotechnology Information. PubChem BioAssay Database. 2017.

36 Samuel Lampa, Jonathan Alvarsson, and Ola Spjuth. SciLuigi Case study workflow - GitHub. https://github.com/pharmbio/bioimg-sciluigi-casestudy. [Online; Accessed 30-July-2018].

37 Science for Life Laboratory. Sarek - an open-source analysis pipeline to detect germline or somatic variants from whole genome sequencing. http://opensource.scilifelab.se/projects/sarek/, (Accessed: 2018/06/01).

38 Heng Li, Bob Handsaker, Alec Wysoker, Tim Fennell, Jue Ruan, Nils Homer, Gabor Marth, Goncalo Abecasis, Richard Durbin, and 1000 Genome Project Data Processing Subgroup. The sequence alignment/map format and samtools. Bioinformatics, 25(16):2078–2079, 2009.

39 Broad Institute. Picard tools. http://broadinstitute.github.io/picard/, (Accessed: 2018/06/01).

40 Aaron McKenna, Matthew Hanna, Eric Banks, Andrey Sivachenko, Kristian Cibulskis, Andrew Kernytsky, Kiran Garimella, David Altshuler, Stacey Gabriel, Mark Daly, and Mark A DePristo. The Genome Analysis Toolkit: a MapReduce framework for analyzing next-generation DNA sequencing data. Genome Res., 20(9):1297–1303, September 2010.

41 Simon Andrews. Fastqc - a quality control tool for high throughput sequence data. https://www.bioinformatics.babraham.ac.uk/projects/fastqc/, (Accessed: 2018/06/01).

42 Alexander Dobin, Carrie A. Davis, Felix Schlesinger, Jorg Drenkow, Chris Zaleski, Sonali Jha, Philippe Batut, Mark Chaisson, and Thomas R. Gingeras. Star: ultrafast universal rna-seq aligner. Bioinformatics, 29(1):15–21, 2013.

43 Konstantin Okonechnikov, Ana Conesa, and Fernando García-Alcalde. Qualimap 2: advanced multi-sample quality control for high-throughput sequencing data. Bioinformatics, 32(2):292–294, 2016.

44 Yang Liao, Gordon K. Smyth, and Wei Shi. featurecounts: an efficient general purpose program for assigning sequence reads to genomic features. Bioinformatics, 30(7):923–930, 2014.

45 Philip Ewels, Måns Magnusson, Sverker Lundin, and Max Käller. Multiqc: summarize analysis results for multiple tools and samples in a single report. Bioinformatics, 32(19):3047–3048, 2016.

46 Go development team. Go FAQ: History of the project. https://golang.org/doc/faq#history.

47 Go development team. Go FAQ: Performance. https://golang.org/doc/faq#Performance.

48 J Paul Morrison. Flow-Based Programming: A new approach to application development. Self-published via CreateSpace, Charleston, 2nd edition, May 2010.

49 Vladimir Sibirov. GoFlow source code repository at GitHub. https://github.com/trustmaster/goflow. [Online; Accessed 16-July-2018].

